# Drowning a frog respiratory rhythm generator in a wash of excitation: State-dependent architecture of a ventilatory oscillator

**DOI:** 10.64898/2026.01.07.698180

**Authors:** Mufaddal I. Baghdadwala, Marina R. Sartori, Richard. J. A. Wilson

## Abstract

Breathing begins at birth and continues to death, providing a unique system to understand the architecture of rhythm-generating circuits in vertebrates. In mammals and frogs, three discrete rhythmogenic brainstem nuclei appear essential for generating different phases of breathing, with individual neurons intrinsically tuned to each phase. This discrete multi-oscillator architecture differs fundamentally from the distributed spinal locomotor networks, where rhythmogenesis spans multiple segments and the active phase of individual neurons depends on network state. However, whether these discrete respiratory oscillators are strictly essential and whether the phase of their neurons remains constant across states is coming under increasing scrutiny.

Here, we focus on the frog buccal oscillator, which shares anatomical and functional similarities with the mammalian preBötzinger Complex. We developed a technique to silence motor neurons in frogs and mapped premotor unit activity within and surrounding the putative buccal oscillator under standard conditions and during heightened network excitability. We found that increasing network excitation caused the rhythmically active area to expand, in part by converting premotor lung units into buccal units.

Next, we located the buccal area using excitatory and inhibitory microinjections and established the minimum number of inhibitory injections required to suppress buccal bursts. We then increased network excitability and again tested whether inhibitory microinjections could suppress the rhythm. While inhibitory microinjections could reverse local excitatory effects, they were unable to abolish the buccal rhythm under elevated excitation.

Our data shows that not only does the region containing rhythmic premotor buccal units enlarge as network excitation increases, but the essential requirement for a discrete, rudimentary buccal oscillator is also lost. Thus, although rostral-caudal segments of the frog brainstem generally contribute to discrete phases of ventilatory motor patterns, our findings suggest that oscillator identity should be refined: oscillators should be viewed as promiscuous flexible functional entities that expand or contract to occupy different extents of the brainstem, rather than as fixed anatomical nuclei.

**Key points summary:**

- The respiratory rhythm is hypothesized to be generated by specialized brainstem circuits comprised of individual non-redundant, discrete, and intrinsically rhythmic oscillators.
- These oscillators are essential in the impoverished network, often generating rhythmic output in isolation. The inspiratory preBötzinger Complex (preBötC) is the most important of these oscillators in mammals.
- Isolated brainstem preparations in frogs have revealed a ventilatory rhythm-generating circuit closely resembling that of mammals. Particularly, the Buccal Area shares anatomical and functional similarities to the preBötC.
- We demonstrate that the discrete Buccal Area is rearranged with mild increases in network excitation to a diffuse network that appears to generate the buccal rhythm instead.
- These findings support the broader hypothesis that respiratory rhythm-generating circuits can switch to being diffuse and redundant, with discrete oscillators quickly drowning in a sea of excitation.

## Introduction

Breathing is a fundamental physiological process. A centrally located neuronal circuit produces the rhythmic, multiphasic patterns of respiratory muscle activation required for normal breathing (Feldman et al., 2013). Defining the architecture of this rhythm-generating circuit across behavioral and experimental states has proved difficult, and the structure and output of the network appear to depend strongly on the state of the preparation being studied (Del Negro et al., 2002a; Smith et al., 1991, 2007; St-John, 2008). In contrast to many vertebrate and invertebrate motor systems where rhythmogenesis is distributed and the active phase of neurons is dependent on state (Fouad et al., 2018; McMahon et al., 2023; Milla-Cruz et al., 2025), the core respiratory rhythm generator for normal breathing is still generally considered to be static with little degeneracy and/or flexibility of neuronal firing (Feldman and Kam, 2015; Gourévitch and Mellen, 2014).

Previously, we speculated that when respiratory networks are isolated from the intact animal or examined in impoverished states, discrete nuclei with a high intrinsic propensity for rhythm generation become unmasked (Wilson et al., 2006). This idea is supported by isolated or simplified brainstem preparations which have revealed multiple endogenously rhythmic oscillators in lamprey, frogs, reptiles, birds and rodents, some of which are only expressed in conditions that activate chemoreceptors (Anderson and Ramirez, 2017; Bongianni et al., 2016; Fortin et al., 1995; Johnson et al., 2016).

In frogs, the isolated brainstem produces coordinated buccal and lung rhythms that closely resemble those *in vivo* (Gdovin et al., 1999; Perry et al., 1995; Sakakibara, 1984). Rhythmic buccal bursts appear to be the default mode, with increasing recruitment of lung bursts when chemoreceptors are activated (Taylor et al., 2003; Torgerson et al., 1997a). Extensive mechanistic studies of this system led us to propose a coupled oscillator architecture in which the rhythmic output of three seemingly discrete loci (the Lung Powerstroke, Lung Priming and Buccal areas) are bound into a single cohesive multiphase motor output (Baghdadwala et al., 2015, 2016).

In mammals, studies point to a comparable organizational framework with inspiration, post-inspiration, and active expiration assigned to spatially distinct rhythmogenic regions, the preBötC, PiCo, and pFRG (Anderson et al., 2016; Onimaru et al., 2009; Smith et al., 1991; Thoby-Brisson et al., 2009). Of these functional nuclei, the most studied is the preBötC which is in the same rhombomere as the frog’s Buccal area (Ballanyi and Ruangkittisakul, 2009; Smith et al., 1991). Like the Buccal Oscillator, activity of the PreBötC (and possibly the PiCO) appears to be the default mode with activity accompanying all overt inspiratory events (Alsahafi et al., 2015). In contrast, and like the frog’s Lung Oscillator, the pFRG is recruited with elevated CO_2_ and/or metabolic rate (Okada et al., 2007; Onimaru et al., 2009; Pisanski and Pagliardini, 2019).

However, while the epicenters of these functional nuclei has become well defined, the apparent size, boundaries, and necessity of these seemingly discrete oscillators are likely state dependent (St-John et al., 2009; Ruangkittisakul et al., 2014; Feldman and Kam, 2015; Sherman et al., 2015; Baertsch et al., 2019). Baertsch et al. (2019) showed that the region capable of producing inspiratory bursts in mice expands when inhibition is reduced and contracts when inhibition is restored. Using *in vitro* and *in vivo* preparations, Baertsch et al. (2019) revealed spatially stochastic origins of inspiratory bursts across a broad region of the ventral medulla and a marked expansion of the active region under conditions of reduced inhibitory tone. These findings support the view that inspiratory rhythmogenesis in rodents is not confined to a single tightly delimited nucleus but is expressed across a flexible and dynamically reconfigurable network.

This dynamic framework aligns closely with our earlier prediction that discrete respiratory oscillators become more sharply defined in hypo-excited networks and less discrete as network excitability increases (Wilson et al., 2006). Importantly, the evidence for such dynamic redistribution of rhythmogenic capacity has so far been limited almost entirely to rodents. Whether this phenomenon represents a general design principle of vertebrate respiratory circuits or a specialization of rodent inspiratory networks remains unknown.

The frog buccal oscillator offers an opportunity to address this gap. The Buccal oscillator and preBötC are likely orthologous rhythmogenic structures derived from the same embryonic rhombomere (Baghdadwala et al., 2015) and likely participate in every ventilatory event (Baghdadwala et al., 2016; Burggren and Doyle, 1986; Wilson et al., 2002). Moreover, the isolated frog brainstem is well oxygenated and produces rhythms that closely match the intact system (Kogo et al., 1994; Okada et al., 1993; Torgerson et al., 1997b), and the buccal oscillator occupies a small and discrete footprint under baseline conditions (Wilson et al., 2002). These features make frogs an ideal comparative system for determining whether state-dependent redistribution of rhythmogenic capacity is likely a conserved vertebrate property.

Here, we test this idea by manipulating network excitability in the isolated bullfrog brainstem and assessing the spatial footprint and functional relevance of the Buccal oscillator. If rhythmogenesis is genuinely dynamic, increased excitation should diminish the functional importance of the anatomically discrete buccal oscillator and enlarge the region capable of generating buccal-like activity. Such a pattern would parallel the dynamic expansion observed in rodents, although driven here by increased excitation rather than reduced inhibition. Our findings support this prediction and indicate that state-dependent redistribution of rhythmogenic capacity is likely a broadly conserved feature of vertebrate respiratory networks.

## Methods

### Ethics

All experiments were performed on juvenile American Bullfrogs, *Rana catesbeiana* (stages 15-24) (Taylor and Kollros, 1946). Animals were obtained commercially from Island Bullfrogs, Nanaimo, BC, Canada. Animal upkeep and experimental protocols were in accordance with the Canadian Council of Animal Care and approved by the University of Calgary Animal Care Committee (protocol AC16-0168).

### Isolated frog brainstem preparation

Animals were anesthetized in MS-222 (0.6 g/l) (Hedrick and Winmill, 2003) until non-responsive to leg or tail pinch. A transverse cut was made at the level of the eyes to expose the cranial cavity followed by transection of the body rostral to hindlimb exit. The cranium was then exposed, and the animal decerebrated at the optic chiasm. A series of longitudinal cuts were made to expose the brainstem and spinal cord. Cranial nerves (CN) were dissected and cut, and a transverse cut was made 1 mm caudal to the hypoglossal nerve root (Spinal Nerve 1; SN1) (Stuesse et al., 1983). The brainstem was then extracted from the cranial cavity and placed in the experimental chamber where the brainstem was pinned down and dura removed. Throughout the surgery, the cranium was superfused with ice-cold artificial cerebrospinal fluid (aCSF; composition in mM: 104 NaCl, 4 KCl, 1.4 MgCl_2_, 10 D-glucose, 25 NaHCO_3_ and 2.4 CaCl_2_) equilibrated with 5% CO_2_ and 95% O_2_. All preparations were allowed at least 30 minutes to stabilize after dissection. Motor output was recorded from CN5, CN7, CN10 and SN1 roots using extracellular glass suction electrodes pulled to have inner tip diameters equal to the diameter of the nerve (**Fig. 1**). Signals were amplified (×10,000) and filtered (300 Hz to 1 kHz) using differential AC amplifiers (model 1700, A-M Systems Inc., Everett, WA, USA) and rectifier and integrated using a moving averager (time constant: 50 ms; MA 821, CWE, Inc., Ardmore, PA, USA). The signals were digitized at 20 kHz and saved as computer files (Axon Digidata board and Axoscope software, Molecular Devices, CA, USA).

**Figure 1:**
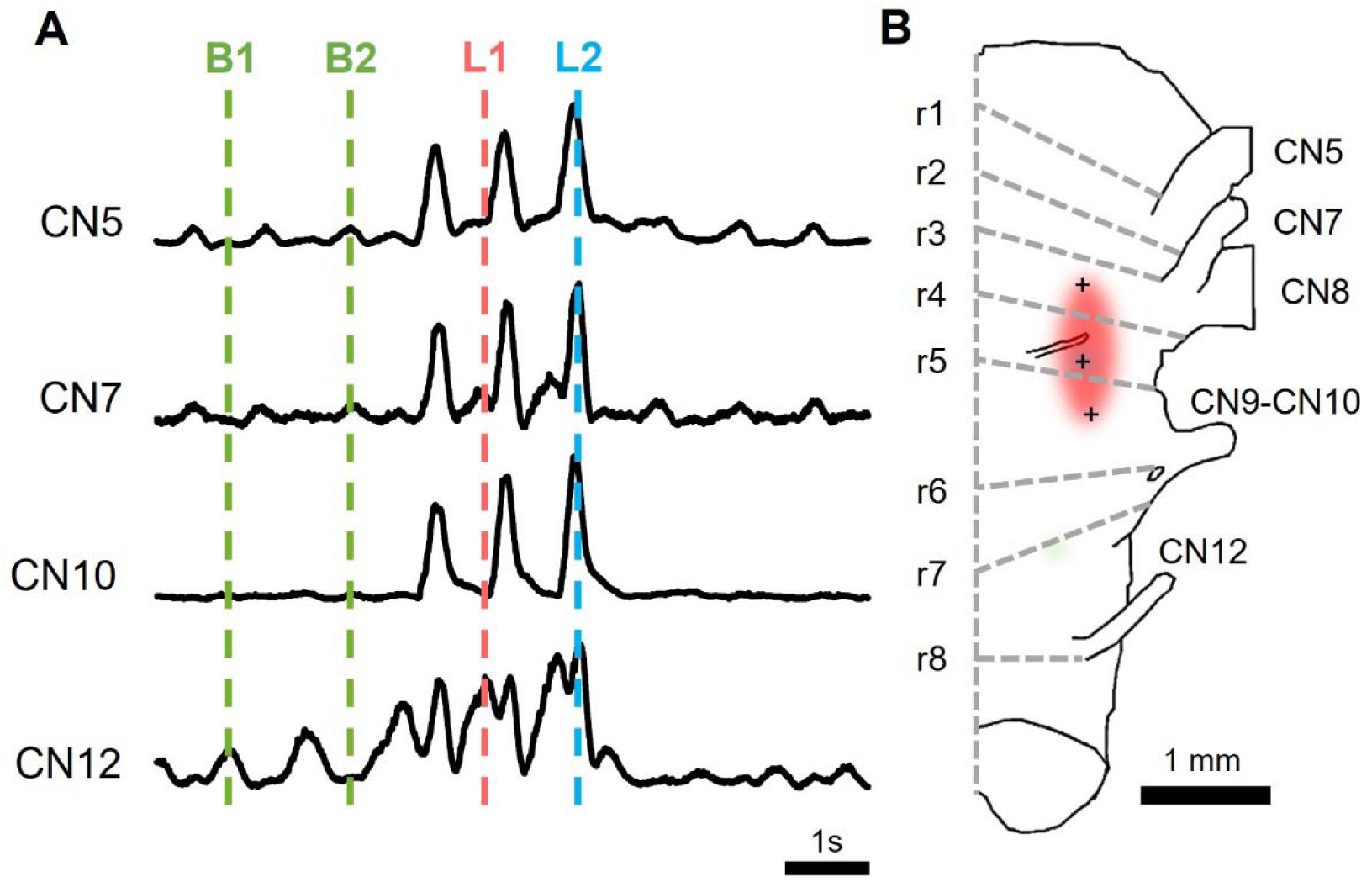
Ventilatory phases of the Bullfrog. **(A)** Extracellular recordings from cranial nerves (CN5, CN7, CN10 and SN1) displaying four phases of ventilation: B1 buccal-depression, B2 buccal-elevation, L1 lung-priming, and L2 lung-powerstroke. **(B)** Anatomic brainstem regions implicated in generating each ventilatory phase.

### General transection and hemisection procedure

After removing the brainstem from the cranium, it was placed in a dish containing high-Mg^2+^ aCSF which was equilibrated with 5% CO_2_ and 95% O_2_. The brainstem was left in this solution for at least 20 minutes before proceeding, to allow abolishment of all synaptic activity. All cuts were made using a sharp scalpel with attention being paid to ensure that the brainstem was not compressed.

### Motor neuron ablation using nerve stimulation

To suppress any motor neuron activity near the Buccal area, we utilized high-frequency stimulation of the nerves to ablate motor neuron pools via excitotoxicity. The extracellular electrodes hooked onto CN10 and SN1 were attached to a stimulator (Grass S88 Stimulator, Grass Medical Instruments, Quincy, MA, USA). The stimulator delivered [8x10 V] 1 ms pulses at 100-400 Hz to a stimulus isolation unit having a 1x voltage multiplier and capacitance coupling. The nerves were bilaterally stimulated for 30-60 minutes under high-Mg^2+^ aCSF (composition in mM: 71.6 NaCl, 4 KCl, 20 MgCl_2_, 10 D-glucose, 25 NaHCO_3_; osmolality of the high-Mg^2+^ solution is adjusted by substitution of sodium and calcium ions). The high-Mg^2+^ solution reliably blocks all synaptic activity by preventing vesicle release due to lack of Ca^2+^ entry. Nerve activity was checked post-stimulation and the nerves were confirmed to be quiescent (**Fig. 2**). Recordings of buccal and lung busts in an unstimulated nerve (CN5) were used as a test of preparation viability. To demonstrate the overall reliability of this approach for inactivating motor neurons, we pooled data from all preparations subject to nerve stimulation (n=34 for CN5 and CN10; n=21 for SN1) and used a two-way ANOVA to compare nerve activity before and after stimulation. The data are present in **Figure 2**.

**Figure 2:**
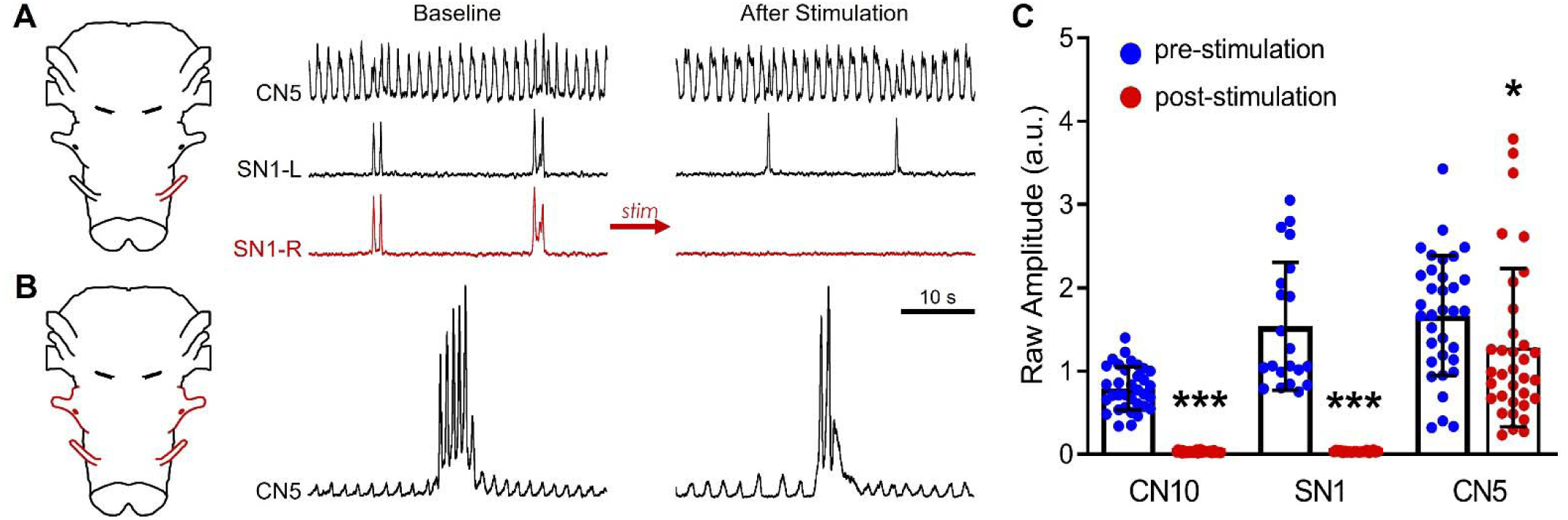
Functional ablation of motor neuron pools to focus on rhythmic interneurons using high-frequency nerve stimulation. **(A)** Extracellular stimulation of SN1-R (red) with high-frequency selectively inhibits ventilatory output in stimulated nerve. **(B)** Bilateral high-frequency stimulation of CN10 and SN1 (highlighted in red) does not affect buccal and lung activity on CN5. **(C)** Quantification of activity (raw amplitude) pre- and post-high frequency stimulation illustrated in Panel B. Bars represent mean+SD (Two-way ANOVA, *** p<0.0001 and * p<0.05, n=34 for CN5 and CN10; n=21 for SN1).

### Buccal Segment dose response

Transverse cuts were made just rostral to the exit of CN10 and caudal to exit of SN1 (**Fig. 3A**). The resultant ‘Buccal segment’ was then placed in the experimental chamber and immobilized using a custom holder (ventral side up). Since the segment seldom produced buccal bursts, 125 nM AMPA was bath applied for 5 min to check the viability of the preparation. Once the preparation was confirmed viable (appearance of bursts on both nerves), a second cut was made to further divide the segment into two. These cuts were either between the nerve exits of CN10 and SN1 (level 1, n=6), or just caudal to CN10 exit (level 2, n=6) (**Fig. 3B-C**). A dose response of bath applied AMPA (60 nM, 125 nM, 250 nM, 500 nM) was then conducted on the two segments to check for bursts.

**Figure 3:**
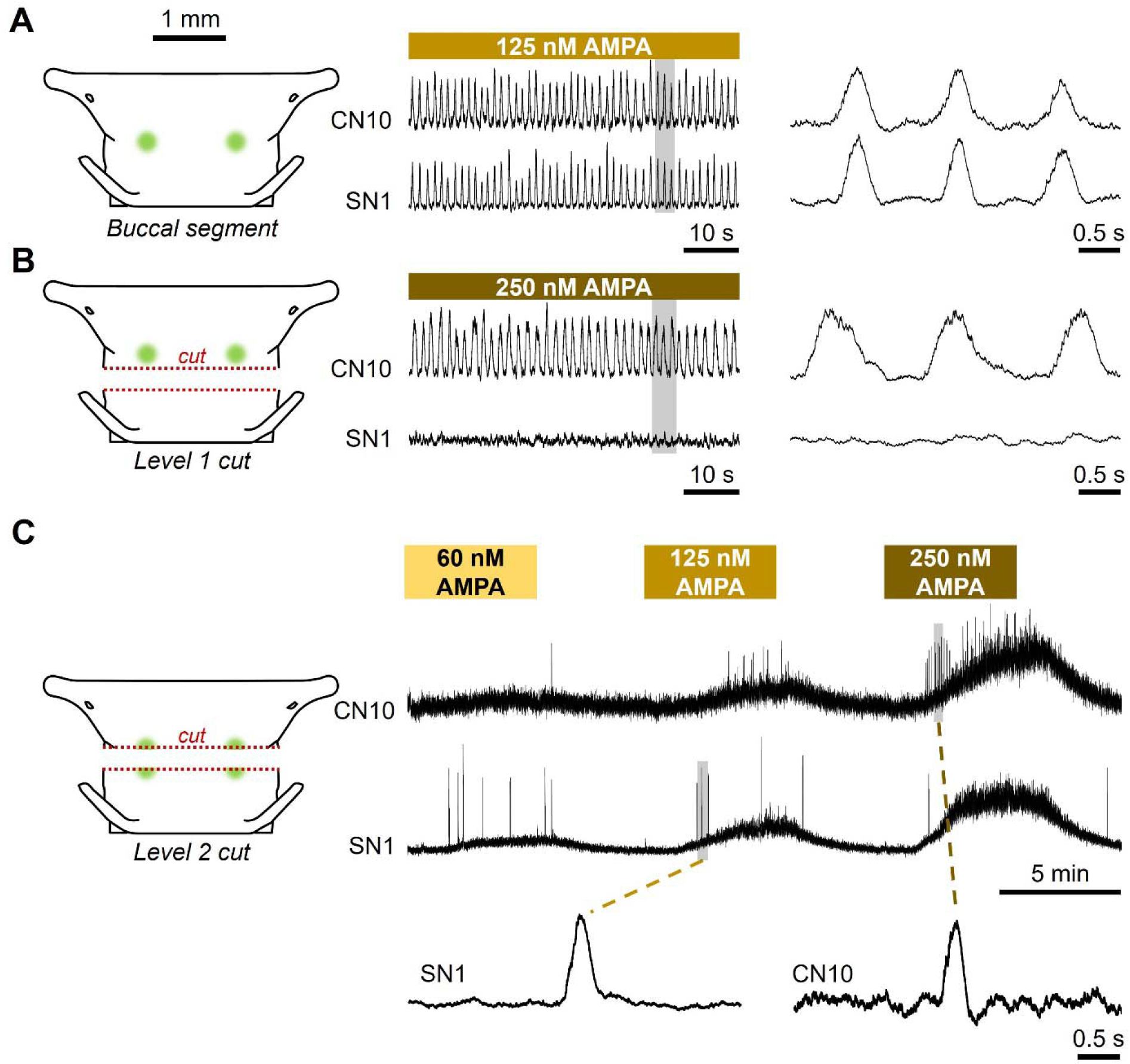
Integrated nerve activity of minimum Buccal Area region capable of producing bursts. **(A)** In this example, a brainstem segments spanning CN10 and SN1 roots produces reliable buccal-like bursts, especially when the network is excited with bath-applied AMPA. **(B)** After transection between CN10 and SN1, just caudal to the previously identified Buccal Area (i.e., *Level 1 cut*; n=6), buccal-like bursts can still be induced with bath-applied AMPA from the rostral slice (containing CN10) but not from the caudal slice (containing SN1). **(C)** After a slightly more rostral cut aimed at transecting the previously identified Buccal Area and yielding a thinner rostral and thicker caudal slice (i.e., *Level 2 cut*; n=6), both slices retain limited capacity to produce bursting with AMPA. Note transient activity with AMPA application, suggesting an optimal range of network excitation required for inducing bursting.

#### Analysis

The number of bursts were counted during each bout (baseline, initial 125 nM AMPA, post-cut 60, 125, 250 and 500 nM AMPA) and the values from each bout were compared to zero using a One-sample t-test. Dose response differences were tested with Friedman test plus Dunn’s multiple comparisons (**Fig. 4**).

**Figure 4:**
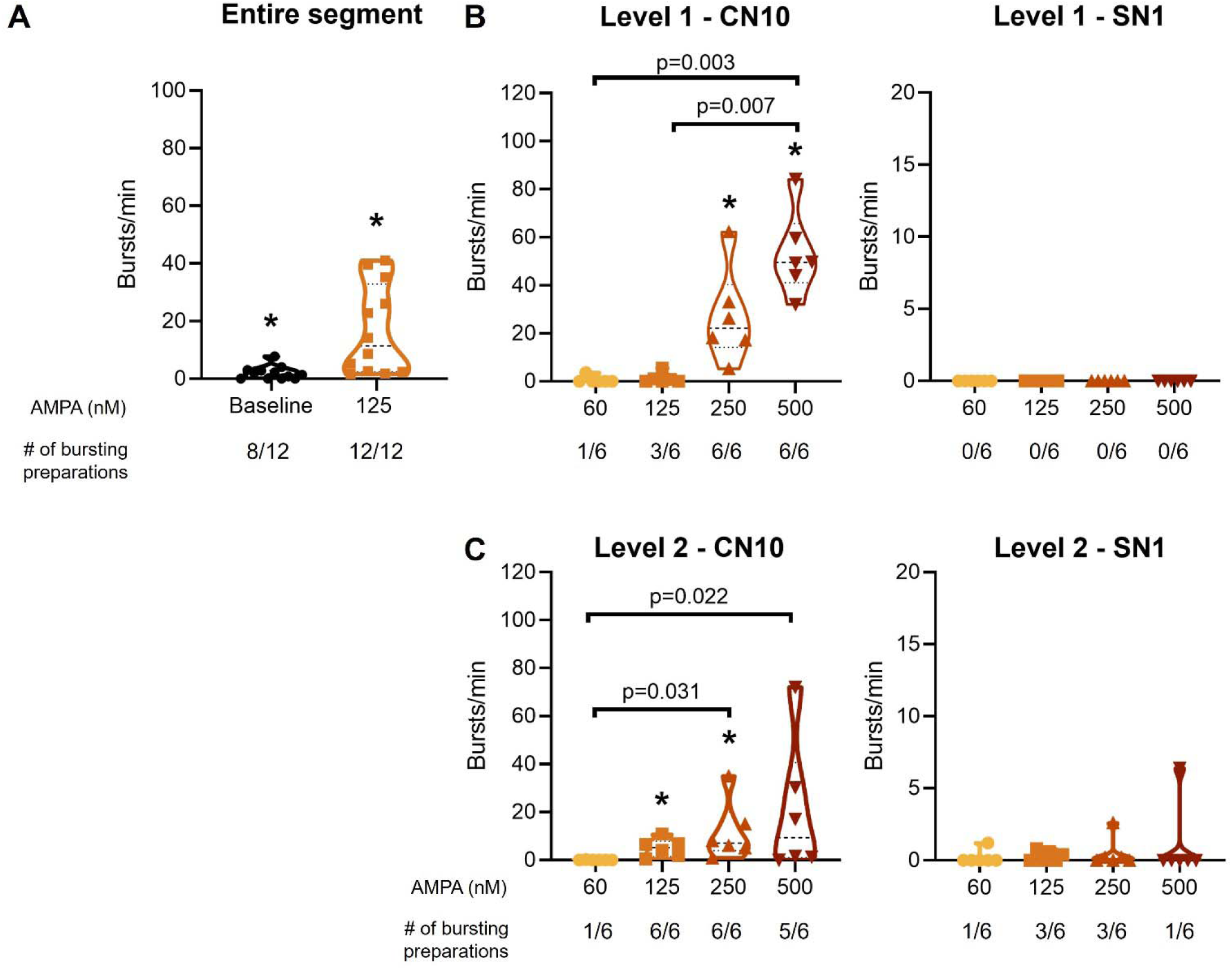
Group data of bursting activity following the transections illustrated in. Fig. 3**. (A)** The isolated buccal segment preparation that includes CN10 and SN1 maintains the capacity to produce bursts both at baseline (8 out 12 preparations) and with 125 nM AMPA (all preparations) (n=12). **(B, C)** Bursting in rostral CN10 slices (*left panels*) and caudal SN1 slices (*right panels*) resulting from the Level 1 cuts (n=6) and Level 2 cuts (n=6), respectively. While neither rostral (CN10) nor caudal (SN1) slices were able to generate spontaneous bursts without AMPA, AMPA caused a significant recruitment of bursting in rostral slices showing persistent bursting capacity in all preparations (compared to zero: p=0.031). In contrast, bursting in caudal slices was practically absent; AMPA failed to induce any bursts in thinner caudal slices, and only induced bursts in 3 of 6 thicker caudal slices which was not enough to reach significance across the group (compared to zero: p>0.25). Bars represent mean±SD. Asterisk symbols denote difference from zero (Wilcoxon one sample test); p values denote dose-response differences (Friedman test plus Dunn’s multiple comparisons).

### Extracellular unit survey, whole brainstem preparation

Following brainstem isolation, CN10 and SN1 were bilaterally stimulated (as described above) to eliminate any motor neuron activity near the Buccal area. Borosilicate glass capillaries (filamented, 1.5mm outer diameter, 1.12mm inner diameter) were pulled using a micropipette puller (Model P-97, Sutter Instruments Co. USA). The electrodes were filled with saturated NaCl solution (tip diameter of ∼10 μm). A systematic extracellular survey was conducted to search for the caudal brainstem which contains the Buccal area (Wilson et al., 2002) for premotor units.

This was achieved using a boxed search between CN10 and SN1 nerve exits with x and y coordinates based on anatomical landmarks on the ventral surface (**Fig. 5A**). Each box having a volume of ∼0.15 mm^3^ was surveyed for 5-10 minutes. Boxes were searched in a pseudo-random order with balanced design. All stable single units detected were documented regardless of their firing pattern with the aim of obtaining over 100 units per condition. Separate experiments were conducted under baseline conditions (normal aCSF, with data pooled from 15 preparations) and under bath application of 60 nM AMPA (with data pooled from 6 preparations). In a subset of six preparations, the brainstem was isolated and CN10 and SN1 were stimulated as described above to suppress buccal motor neuron activity. The brainstem was then transected rostral to the exit of CN10 and caudal to the exit of SN1, functionally and anatomically isolating the buccal region from the lung region. A similar box search was used as above to identify units. Separate experiments were conducted under baseline conditions (normal aCSF) and under bath application of 60 nM AMPA.

**Figure 5:**
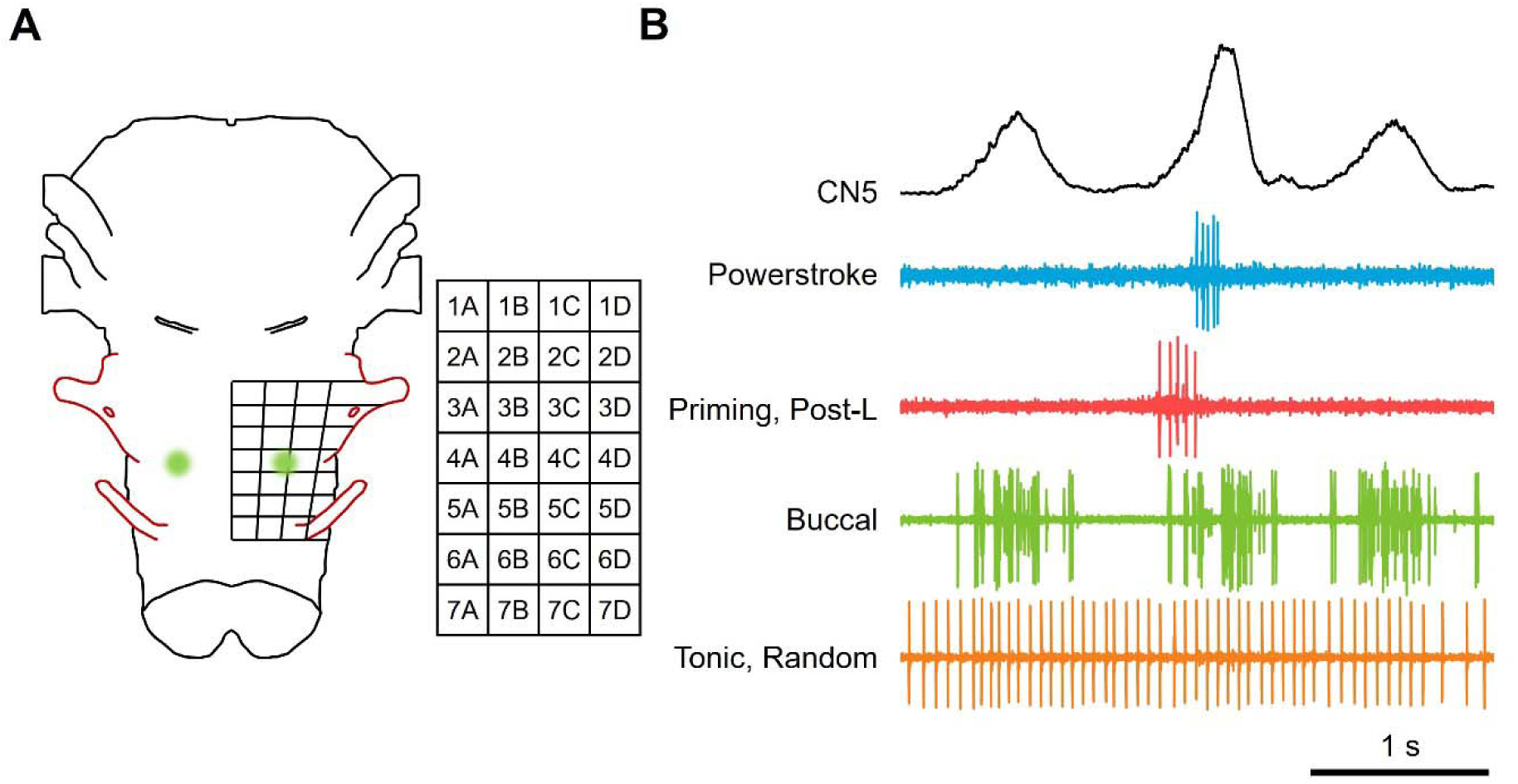
Survey of rhythmic units in the buccal region. **(A)** The buccal region was divided into 28 boxes based on a grid defined using anatomical landmarks to facilitate a systematic survey of units in the buccal region. **(B)** Single units were defined according to their patterning of activity. Subsequent figures illustrate the number of units identified in each anatomically defined box.

#### Analysis

Units were pooled from multiple preparations. Units were sorted by comparing their firing patterns to the fictive motor output recorded from CN5 (**Fig. 5B**). Four categories were used: (1) powerstroke, (2) priming/post-lung, (3) buccal, (4) tonic/random. Units that fired only during lung bursts in CN5 were considered powerstroke units. Units that fired before or after lung bursts in CN5 were considered priming/post-lung. Any units that fired during buccal bursts as well as lung bursts in CN5 were labeled buccal units (no units were found which fired only during buccal bursts in CN5). Finally, units with activity uncorrelated to bursts in CN5 were labelled tonic/random (**Fig. 5B**). A chi-squared test was done to determine statistical differences in the proportion of units of each category in experiments with and without 60 nM AMPA. The subset of six preparations in which the brainstem was cut just rostral to the exit of CN10 and caudal to exit of SN1 lacked functional nerve output. To analyze data from these preparations, units were sorted into 1 of 2 categories: (1) rhythmic bursting and (2) tonic/random. As the powerstroke and priming burst-generating areas were absent, the rhythmic units were assumed to be buccal. A chi-squared test was used to test for significant differences in the proportion of units in experiments with and without 60 nM AMPA.

### Microinjections

Hemisected preparations were utilized for the microinjection experiments. Following isolation of the brainstem, a longitudinal cut along the midline was made and the hemisected brainstem was immobilized using pins in the experimental chamber (ventral side up). Dual or 4-barrel glass capillary tubes (filamented, 1.2 mm outer diameter, 0.6 mm inner diameter) were heat-pulled into micropipettes and back filled with AMPA (5 µM) and GABA (5 mM) in separate barrels. The open end of the micropipettes was attached to a multi-channel Picospritzer (General Valve Corporation, Fairfield, NJ, USA) which was used for pressure-controlled microinjections; the volume of the injection was controlled by adjusting the duration (in milliseconds) of the pressure valve being open. To calculate the volume of each injection, a large cumulative injection (40 injections of 1 s each) was carried out at the end of each experiment to cause a measurable change in the meniscus of the injecting capillary; the change in level of meniscus along with the internal diameter of the capillary was used to calculate the volume of the large injection (volume = πr^2^h; r: internal radius of capillary tube, h: change in level of meniscus) which allowed us to estimate the volume of each small injection. An area was found where small injections of 5 µM AMPA (5±1 nl) and 5 mM GABA (14±4 nl) could reset and inhibit the buccal rhythm, respectively. This identification protocol is similar to the one utilized by Wilson et al. (2002) to first identify the Buccal area. Once the buccal area was identified, three sets of injections were conducted: 1) AMPA-only injections, 2) GABA-only injections, 3) AMPA-GABA injections. The AMPA-GABA injections entailed injection of AMPA and then 3-5 seconds later, injection of GABA; these were done to ensure that the volume of 5 mM GABA was sufficient to completely overcome the effects of the preceding 5 µM AMPA injection (**Fig. 9A**). Each set of injections was repeated at least 3 times in a given preparation, and responses for each repeat set averaged during analysis to yield one value per animal. At least 3 minutes of washout was provided between each set of injections. After the completion of this protocol, 60 nM of AMPA was bath applied and the entire protocol was repeated (**Fig. 9B**). The placement of the electrode as well as the injection volumes were kept constant between baseline and 60 nM AMPA treatments.

#### Analysis

The intervals between subsequent buccal bursts (inter buccal interval, IBI, in seconds) immediately before and after injections were measured and used to calculate the instantaneous frequency (IF; bursts/seconds):

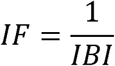

To assess effects of injections on burst activity, we tested the fractional change in instantaneous frequency (a) using a 2-way repeated measures ANOVA (Factors: injection type and bath application) and (b) comparing each condition against zero (indicating no change) using one-sample t-Tests:

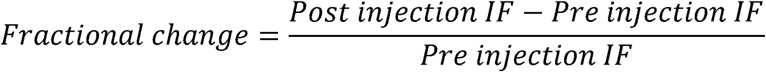

### Simultaneous unit recordings and microinjections

We used whole brainstem preparations which had undergone the CN10 and SN1 motor neuron ablation protocol for these experiments. Simultaneous single-unit recordings and focal microinjections were performed with a five-barrel micropipette. Pipettes were heat-pulled and the tips broken to yield final diameters of 15-20 µm. The central barrel was filled with 150 mM NaCl and connected to a differential amplifier via a silver wire electrode for extracellular unit recording. The remaining four barrels were filled with 5 µM AMPA and connected to a four-channel pico-spritzer that generated a digital pulse during injection. Digital pulses were recorded along with the electrophysiological signals to indicate the time of AMPA injections. Ventilatory motor patterns were monitored from CN5 using a glass suction electrode connected to a differential amplifier.

Injection sites were first identified within the buccal region as locations at which brief AMPA application produced a qualitative increase in buccal burst activity. Injection pressure and pulse duration were adjusted to determine the minimum stimulus required to reliably enhance buccal bursting, thereby minimizing drug spread. The micropipette was then repositioned within a ∼50 µm³ volume to identify a nearby lung-associated unit, defined operationally as a neuron active during lung bursts but silent during buccal bursts. AMPA was subsequently injected at this site, and when stable unit isolation was maintained, the effects on lung-associated neuronal activity were assessed. Neuronal signals were amplified (×10,000), band-pass filtered (300 Hz–1 kHz), digitized at 20 kHz, and stored for offline analysis.

#### Analysis

While making recordings we targeted prominent single lung units (active during lung bursts and not buccal bursts). However, we also embarked on a formal unit analysis offline to ensure the same unit was recorded before and after AMPA injection. Spike sorting was performed using an automated pipeline implemented in Python. Raw signals were high-pass filtered (300 Hz, 3rd-order Butterworth, zero-phase) and candidate spikes identified using an amplitude threshold of 2.5× the estimated noise level (median absolute deviation / 0.6745) with a minimum refractory period of 3 ms. Spike waveforms (±1.5 ms around each peak) were extracted and subjected to principal component analysis. Clustering was performed on the first two principal components combined with peak amplitude using k-means clustering. The optimal number of clusters (1–4) was determined by maximizing the silhouette score, with a minimum threshold of 0.10 required for multi-unit classification. Units were relabeled in ascending order of mean PC1 value for consistency across preparations. To identify respiratory bursts, the CN5 signal was low pass filtered (2 Hz cutoff, 4th-order Butterworth, zero-phase). Peaks were identified using prominence-based detection (prominence threshold: 0.2× signal standard deviation) with a minimum inter-burst interval of 0.7 s. Lung and buccal bursts were distinguished by peak amplitude using k-means clustering (k=2) on absolute peak heights. The boundary between lung (larger amplitude) and buccal (smaller amplitude) bursts was set at the buccal cluster center plus 30% of the distance to the lung cluster center. To avoid contamination from excitation caused by transitional activity, the first four buccal bursts following each lung burst were excluded from buccal-phase analyses. Spikes were classified as "lung-associated" or "buccal-associated" based on temporal proximity (within ±0.8 s) to the nearest burst peak. The largest-amplitude unit was identified and designated as the putative "lung unit" based on its firing pattern under baseline conditions prior to AMPA injection (i.e., active during lung bursts but not buccal bursts).

To compare spike timing across bursts of variable duration, spike times were converted to phase coordinates relative to each burst cycle. Phase was defined as follows: phase 0 corresponded to the burst peak; phase −π corresponded to the midpoint between the current burst peak and the preceding burst peak; phase +π corresponded to the midpoint between the current burst peak and the following burst peak. This peak-centered, piecewise-linear mapping normalized burst timing while preserving the relationship between spike timing and burst phase. Phase windows were capped at a maximum duration of 1.6 s to prevent inclusion of spurious inter-burst CN5 activity.

Phase-recruitment histograms were computed by binning spike phases into 15 equally spaced bins spanning −π to +π. To assess whether spike timing was significantly modulated by burst phase, a Z-score was calculated for each bin by comparing the observed spike count to the expected count under a uniform (null) distribution. The expected count per bin was n/15, where n is the total spike count, with standard deviation calculated from the binomial distribution: σ = √(n × p × (1−p)), where p = 1/15. Bins with Z > 1.65 (one-tailed, α = 0.05) were considered to show significant phase-locked recruitment. Phase-recruitment data was obtained from five stable lung units. Phase-recruitment heatmaps were constructed with Z-score profiles across bins, and violin plots showing the distribution of spikes per burst were generated for each unit, with individual burst spike counts plotted as points and kernel density estimates indicating the distribution shape.

## Results

### Buccal-like bursting produced from two independent slices

To test whether the rhythmicity of the buccal oscillator is preserved under extreme reduction, we attempted to reduce the buccal oscillator to a small brainstem slice. We started with the buccal segment preparation spanning from CN10 to SN1 (n=12 total; **Fig. 3A**). Bursting properties are summarized in **Table 1** and **Fig. 4**. 8 out of the 12 preparations produced spontaneous bursts at a frequency of 2.00 ± 2.28 bursts per minute (bpm) (p=0.008, compared to zero; **Fig. 4A**). With bath applied 125 nM AMPA, all 12 preparations exhibited bursting, demonstrating all preparations retained bursting capacity (16.75 ± 15.49 bpm; p<0.001, compared to zero). In the 8 preparations that produced spontaneous bursts, 125 nM AMPA increased the burst frequency from 3.00 ± 2.18 to 23.23 ± 15.12 bpm. Next, we transected the buccal segment into slices as described below.

**Table 1.**
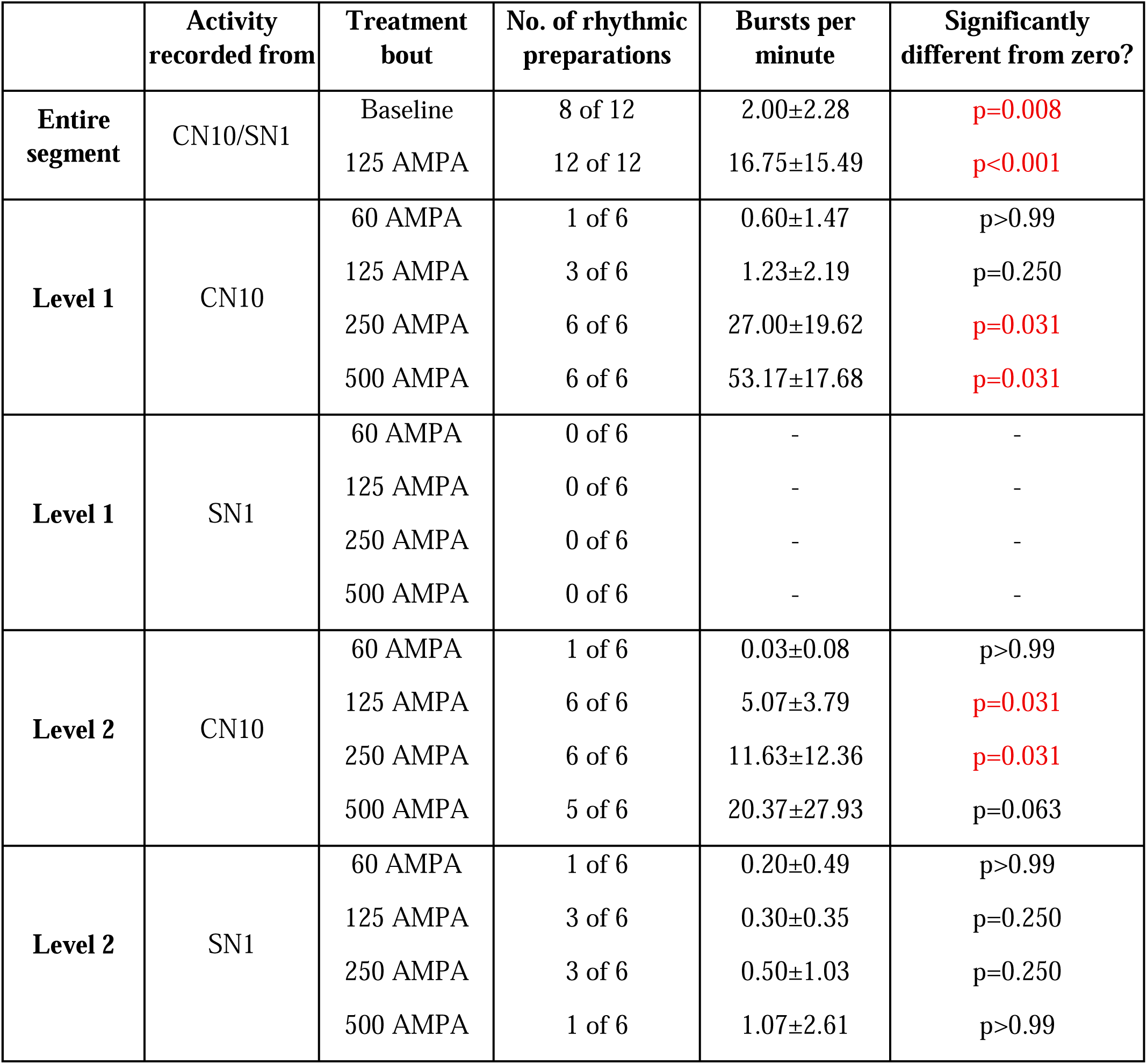
Summary of buccal segment preparations. Number of buccal slices (and their corresponding levels – Level 1 or Level 2) that displayed bursting in response to different concentrations of AMPA (Wilcoxon One sample Test).

#### Level 1 cuts

In half the preparations (n=6) cuts were made equidistant between CN10 and SN1 nerve exits (**Fig. 3B**). Neither rostral nor caudal slice had spontaneous activity. In the rostral slice containing the CN10 (*CN10 slice*), 1 preparation showed activity at 60 nM AMPA (mean 0.6 ± 1.47 bpm; p>0.99), 3 preparations at 125 nM (1.23 ± 2.19 bpm; p=0.250), and all 6 at 250 nM (27.0 ± 19.62 bpm; p=0.031) and 500 nM (53.17±17.68 bpm; p=0.031) (*left panel in* **Fig. 4B**, **Table 1**; p values compared to zero). No bursting activity was found in the caudal slice containing SN1 (*SN1 slice; right panel in* **Fig. 4B**) at any of the AMPA doses.

#### Level 2 cuts

These cuts were made at the caudal edge of CN10 nerve exit (**Fig. 3C**, n=6). Both rostral and caudal slices produced bursting activity at varying levels of AMPA stimulation. For the rostral slice containing CN10, 1 preparation exhibited activity in 60 nM AMPA (0.03 ± 0.08 bpm; p>0.99) all preparations in 125 nM (5.07 ± 3.79 bpm; p=0.031) and 250 nM AMPA (11.63 ± 12.36 bpm; p=0.031), and 5 preparations showed bursting activity at 500 nM AMPA (20.37 ± 27.93 bpm; p=0.063) (*left panel in* **Fig. 4C**; **Table 1**, p values compared to zero). For the caudal slice containing SN1, one preparation exhibited bursting activity at 60 nM AMPA (0.20±0.49 bpm; p>0.99), three at 125 nM AMPA (0.30±0.35 bpm; p=0.025), three at 250 nM AMPA (0.50±1.03 bpm; p=0.025), and one at 500 nM AMPA (1.07±2.61 bpm; p>0.99) (*right panel in* **Fig. 4C**; p values compared to zero).

Note that after slicing the buccal segment preparations at Levels 1 and 2, some slices only responded transiently to AMPA stimulation, producing bursts only when AMPA was being washed in or washed out of the dish. We suspect this is indicative of there being a narrow range of excitation which is conducive to bursting in the slice. This phenomenon is visible in the sample traces in **Fig. 3C**. To account for this type of response, and to prevent any data selection, we counted all bursts occurring during the 5 min AMPA treatment and then averaged the values to produce a burst-per-minute value.

The transection experiments revealed that reducing the buccal oscillator to a slice is possible but only with supplemental AMPA excitation. Also, our ability to produce two slices that can produce bursts is potentially indicative of the entire region having bursting ability when excitation is raised.

### Increased excitation changes the proportion of rhythmic premotor units in the brainstem

The possibility that the entire buccal region has recruitable rhythmic ability upon increased excitation provoked us to investigate how the network architecture changes when AMPA is bath applied. The first set of experiments to test this were conducted on the whole isolated brainstem after CN10 and SN1 motor neuron pools were bilaterally ablated using high-frequency stimulation (**Fig. 2A-C**). We utilized a box grid (**Fig. 5A**) to guide our extracellular electrode placement and to ensure consistency between experiments. Extracellular units were divided into 4 categories (**Fig. 5B**). Two sets of experiments were conducted (using two sets of preparations), one under baseline conditions (n=15) and the other under 60 nM AMPA (n=6). **Figure 6** displays the distribution and density of units under both these conditions. We found a total of 145 units in baseline condition and 207 units with 60 nM AMPA. 40.0% of units were classified as powerstroke with regular aCSF. Significantly fewer powerstroke units were recorded in the presence of 60 nM AMPA (22.2%; Chi-squared test and Bonferroni; p=0.001). Priming/post-L units also significantly decreased from 12.4% with regular aCSF to 1.9% with AMPA (Chi-squared test and Bonferroni; p<0.001). Buccal units increased significantly with AMPA from 9.7% to 22.2% (Chi-squared test and Bonferroni; p=0.008). Tonic/random units also increased significantly from 37.9% to 53.6% with AMPA (Chi-squared test and Bonferroni; p=0.015). A subset of experiments was also conducted on the buccal segment preparation. These followed a very similar protocol to the whole brainstem experiments above. Extracellular units were divided into two categories, putative buccal and tonic; **Figure 7** displays the density and distribution of these units. A total of 110 units pooled from three preparations were found with regular aCSF and a total of 137 units pooled from three additional animals were found with 60 nM AMPA. The proportion of buccal units increased significantly from 6.4% in regular aCSF to 53.3% with AMPA (Chi-squared test and Bonferroni; p<0.001), and a corresponding decrease in the number of tonic/random units from 93.6% to 46.7% (**Fig. 7**).

**Figure 6:**
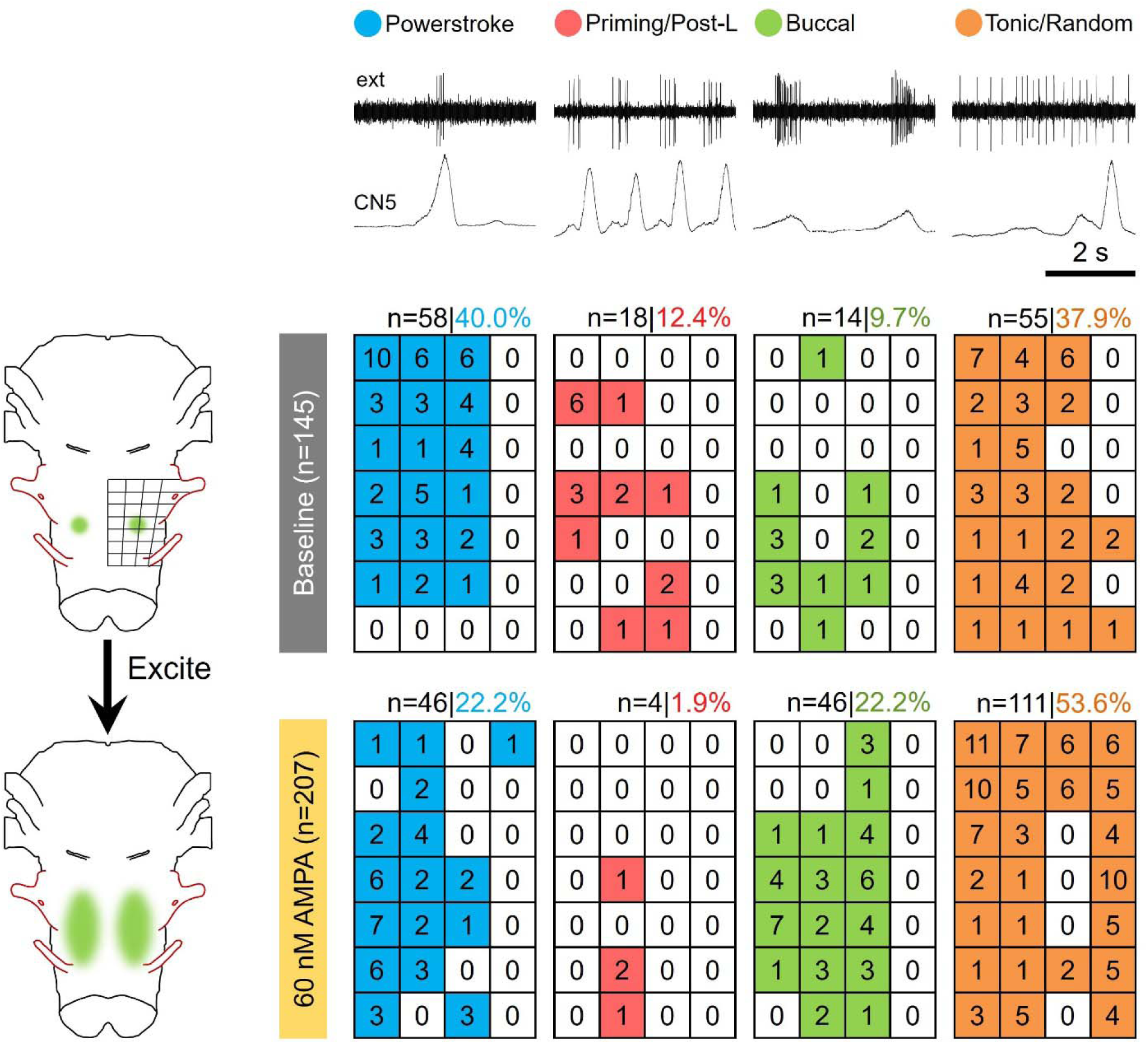
In the isolated intact brainstem, network mild excitation increases the density and distribution of premotor buccal units in the buccal region. CN10 and SN1 (red) motor neurons were functionally ablated before unit surveys. The raster displays the survey results. The numbers are indicative of the frequency of units found in each location from all preparations. Top panel displays unit frequencies under baseline conditions for 145 units pooled from 15 preparations. Bottom panel is from 207 units pooled from 6 preparation in the presence of 60 nM AMPA to cause mild network excitation (Chi-squared test, Bonferroni correction for multiple comparisons; there were significant differences in the proportion of all categories of extracellular units between the two groups; p<0.01). Ext: extracellular unit recording.

**Figure 7:**
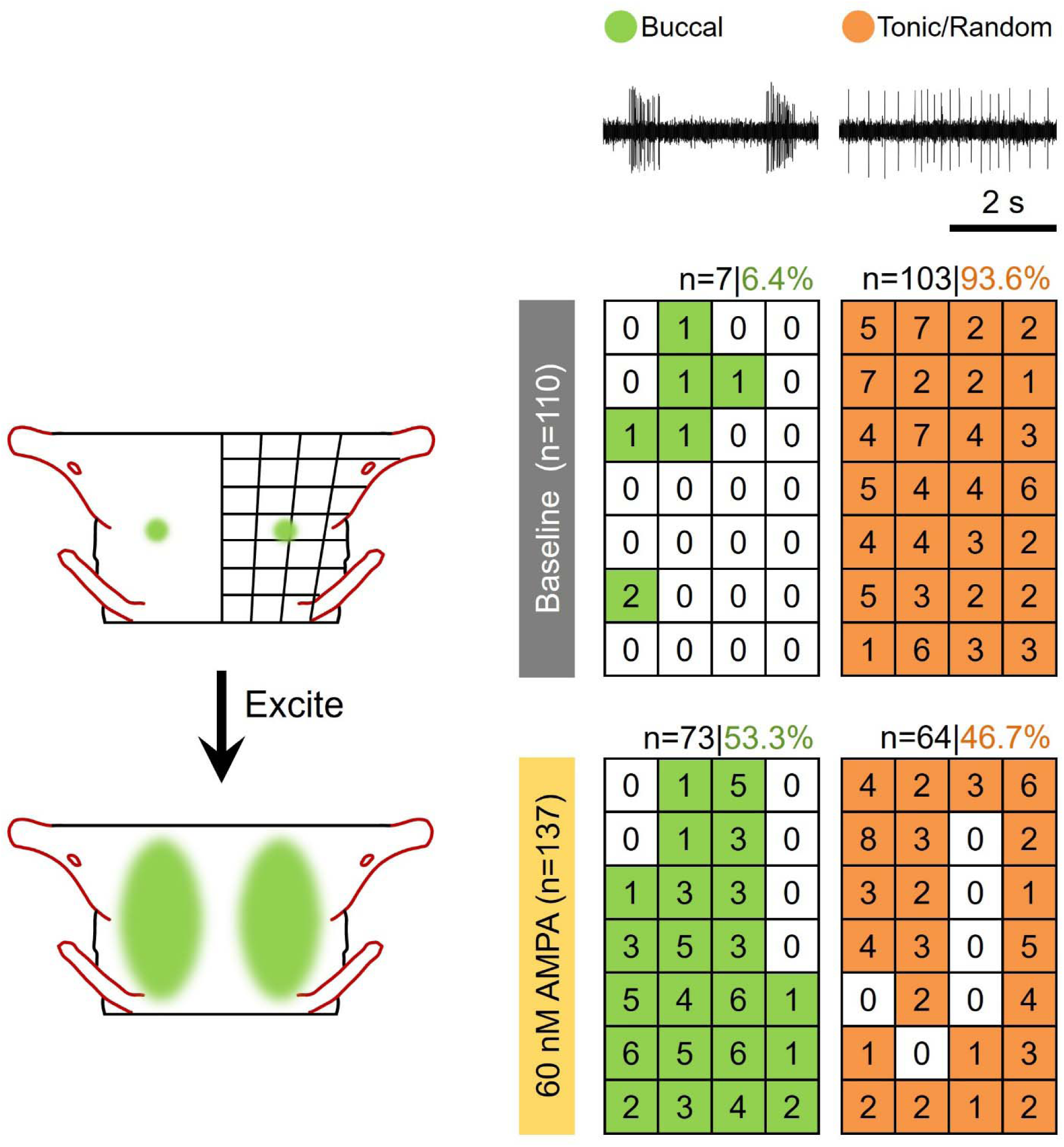
In the buccal segment encapsulating the buccal region, mild network excitation increases the density and distribution of premotor buccal units. CN10 and SN1 (red) motor neurons were functionally ablated before unit surveys. The rasters display the survey results from 247 units pooled from three preparations. The numbers are indicative of the frequency of units found in each location. Top panel displays unit frequencies under baseline conditions from 110 units; bottom panel is from 137 units in the presence of 60 nM AMPA to cause mild network excitation. (Chi-squared test, Bonferroni correction for multiple comparisons; there was a significant difference in the proportion of buccal units between the two groups; p<0.01).

### Lung units can transform into buccal units under excitation

During our analysis of the extracellular survey in the whole brainstem, we observed an overall decrease in the proportion of lung units and a concomitant increase in buccal units during bath application of AMPA (60 nM), suggesting that network excitation alters premotor unit identity. To determine whether these population-level changes were linked at the level of individual neurons, we recorded lung units under baseline conditions and then locally increased excitation by micro injecting ∼0.30 nl of 5 µM AMPA within 10 µm of the recording electrode. We identified and held five lung units (pooled from four animals; location 2B, **Fig. 5A**; 300–400 µm below the dorsal surface) that transformed into buccal units following microinjections. Transformation was confirmed by offline PCA. When post-AMPA buccal bursts were divided into 15 temporal bins, median bin spike counts were significantly greater than zero (one-sample t-test, t = 5.29, p = 0.003; 46.8 ± 17.7) and were non-randomly distributed, with peak bin z-scores exceeding chance levels (z = 1.65; one-sample t-test, t = 3.54, p = 0.012; 3.32 ± 0.94). Together these data suggest a robust induction of buccal-like activity (**Fig. 8**).

**Figure 8:**
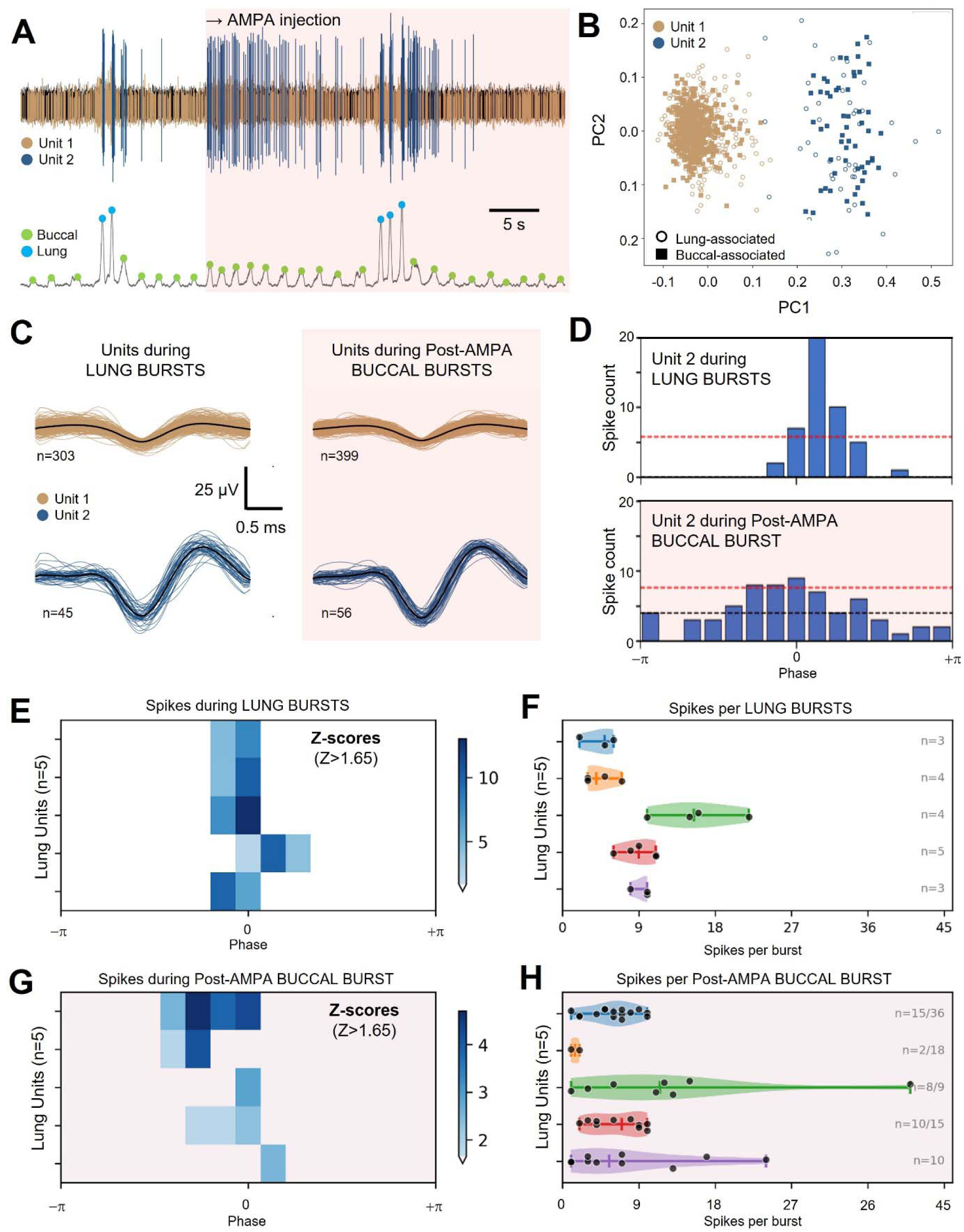
Local network excitation transforms the operational identity of premotor neurons. **(A)** Representative extracellular recording from the caudal brainstem showing activity of two sorted units following local AMPA injection (shaded red). Top trace: raw extracellular signal with spikes color-coded. Bottom trace: integrated CN5 activity with lung bursts (blue circles) and buccal bursts (green circles). Before local AMPA injection, Unit 1 (brown) is active during both buccal associated CN5 bursts, the operational definition of a *buccal unit*. In contrast, Unit 2 (blue) shows lung burst-associated spikes but no buccal burst-associated spikes once effects of a lung episode have subsided, the operational definition of a *lung unit*. However, following AMPA application, this lung unit is transformed into a buccal-lung unit, illustrating local excitation causes network reconfiguration. **(B)** Principal component analysis (PC1 vs PC2) of spike waveforms demonstrating clear separation between the two most prominent units (see Results for silhouette scores and Mahalanobis distances). Marker shape indicates burst association: open circles denote spikes occurring within ±0.8 s of a lung burst peak; filled squares denote buccal-associated spikes. (**C**) Spike-triggered waveform averages for Units 1 and 2 comparing activity during lung bursts (left) versus post-AMPA buccal bursts (right). Individual waveforms shown in brown or blue with mean waveform overlaid in black. The consistent waveforms across conditions confirm unit identity (spike correlation coefficients: r>0.95). **(D)** Phase histograms for Unit 2 (the lung unit) showing spike timing relative to CN5 burst phase (phase 0 = burst peak; ±π = inter-burst midpoints). Top: during lung bursts, Unit 2 fires preferentially near the burst peak. Bottom: during post-AMPA buccal bursts, Unit 2 exhibits a similar phase-locked firing pattern, demonstrating recruitment of this lung unit during buccal activity. Dashed red line indicates the Z-score threshold (Z > 1.65) for significant recruitment above uniform baseline; dashed black line indicates median spike count. **(E, F)** Meta-analysis across five lung units (rows) showing phase-recruitment heatmaps and spikes per burst represented as violin plots during lung bursts, respectively (n=3–5 lung bursts analyzed per unit). Blue color intensity in ***E*** indicates Z-score for recruitment above chance (uniform distribution). Dots in ***F*** indicate individual burst spike counts. **(G, H)** Recruitment of same five lung units during post-AMPA buccal bursts (n= consecutive buccal bursts with lung unit activity / total consecutive buccal bursts available to analyze). The spiking and recruitment pattern during post-AMPA buccal activity in **E-G** is consistent with local excitation causing network reconfiguration.

**Figure 9:**
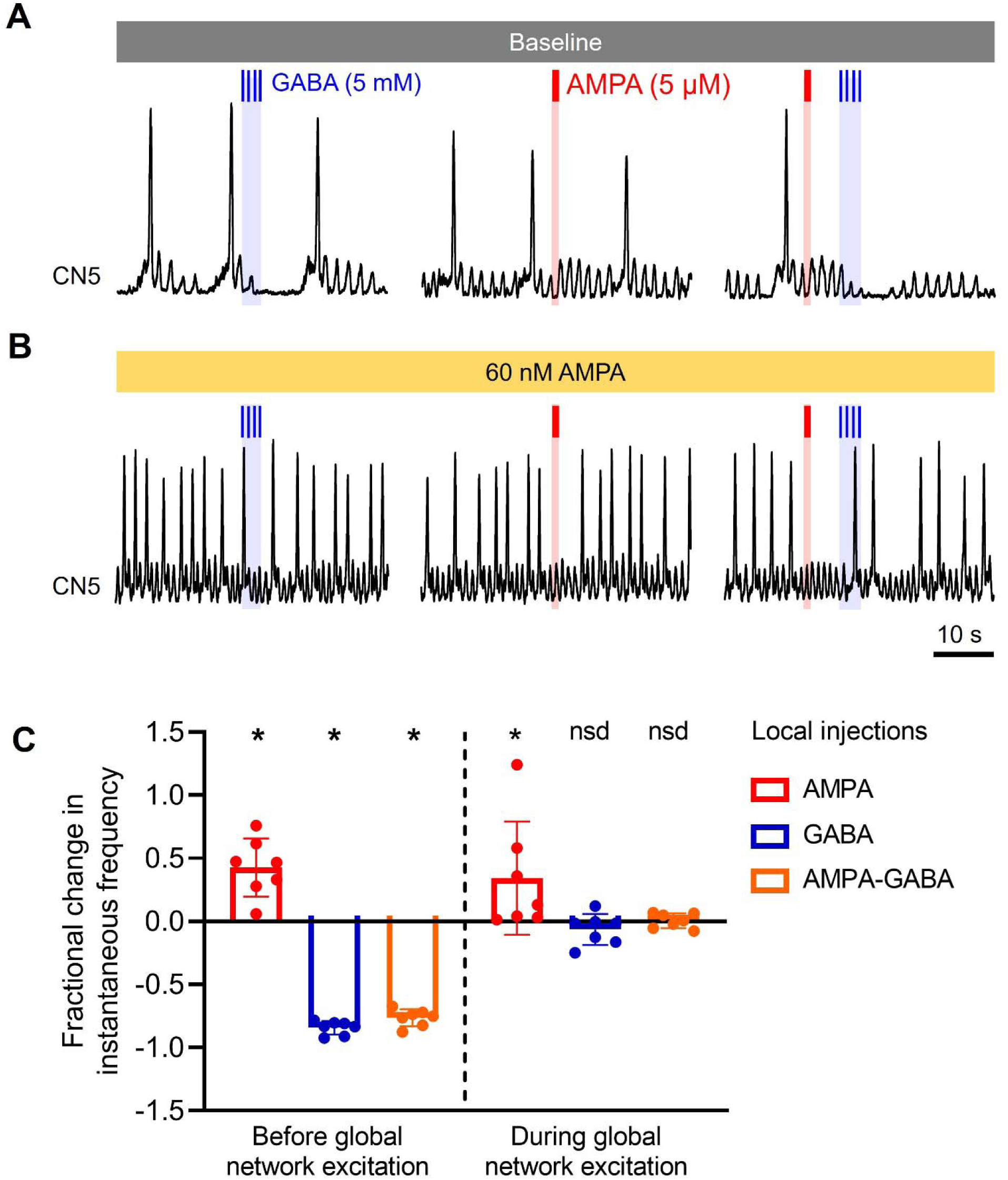
Demonstration that sufficiency and necessity of the Buccal Area is dependent upon the excitation state of the network. **(A)** Effect of AMPA (∼5 nl, 5 µM; sufficiency test) and GABA (∼15 nl, 5 mM; necessity test) microinjections into the Buccal Area of an isolated hemisected brainstem preparation. AMPA causes a brief increase in buccal burst frequency and GABA causes the buccal bursts to momentarily cease; injections of AMPA immediately followed by GABA demonstrate that GABA can completely counteract the excitatory effects of local AMPA injections. **(B)** When the above protocol was repeated with bath application of 60 nM AMPA, the effect of AMPA persist but GABA injections fail to cease buccal bursting suggest the discrete buccal area is replaced by a distributed rhythm generating network. **(C)** Fractional change in instantaneous frequency before and during global network excitation with 60 nM AMPA (One sample t-Test; *\**, p<0.01; *nsd*, no significant difference). Bars represent mean+SD; n=7.

PCA-based spike sorting confirmed unit identity before and after AMPA injection. In all cases, optimal clustering was achieved with k = 2 with one cluster corresponding to pre-AMPA lung units. Clustering yielded strong silhouette scores (median = 0.849, range = 0.661–0.879) and distinct clusters in principal component space, as reflected by Mahalanobis distances (median = 9.08, range = 3.22–11.09). Importantly, waveform correlations between lung unit spikes recorded at baseline and during buccal bursts recorded post-AMPA were consistently high (median r = 0.987, range = 0.957–0.997), providing strong evidence that pre-AMPA lung units were recruited during buccal bursts under conditions of elevated network excitation.

### Buccal area loses importance under increased excitation

The spread of rhythmically active units hinted that the expanded buccal region might have diffuse rhythm generating ability when excited with AMPA. We wanted to test the implications of this increase in rhythmicity on the necessity and sufficiency of the previously discrete buccal oscillator. We utilized a hemisected preparation (to prevent the contralateral buccal area from compensating) and identified the Buccal area using 5 µM AMPA (5±1 nl) and 5 mM GABA (14±4 nl) microinjections. A total of 15 experiments were conducted, but 9 were excluded either because the Buccal area was not successfully found or because the preparation was overdriven with a high frequency of lung bursts and low frequency of buccal bursts. The Buccal area was located at a depth of 257.5±7.5 µm from the dorsal surface and at location 2B on the grid. With brainstem bathed in regular aCSF (**Fig. 9A**), AMPA injections caused increase in instantaneous frequency (from 0.69 ± 0.23 to 1.01 ± 0.48 bursts/sec, leading to a positive fractional change that was significantly different than zero (**Fig. 9C**, One sample t-test compared to zero, p=0.009). GABA, or AMPA followed by GABA injections under baseline conditions reliably caused cessation of buccal bursts consequently reducing instantaneous frequency (GABA: from 0.64 ± 0.11 to 0.09 ± 0.04 bursts/sec; AMPA-GABA: from 0.71 ± 0.11 to 0.16 ± 0.04 bursts/sec), leading to a significant negative fractional change (**Fig. 9C**, one sample t-test compared to zero, p<0.001 for both GABA and AMPA-GABA injections). When 60 nM AMPA was bath applied (**Fig. 9B**), changes caused by the same injected volume of AMPA (from 0.92 ± 0.30 to 1.22 ± 0.38 beats/sec), GABA (from 1.03 ± 0.13 to 0.93±0.12 bursts/sec) or AMPA-GABA (from 1.02 ± 0.14 to 1.04 ± 0.15 bursts/sec) caused no significant change in fractional instantaneous frequency (**Fig. 9C**, one sample t-test compared to zero, p>0.05). Thus, when the network is globally excited using a low concentration of AMPA (60 nM), the ability of a locally applied AMPA concentration that is two orders of magnitude higher (5 µM) to selectively stimulate the buccal area is lost, and GABA is similarly unable to terminate buccal bursting. This indicates that under conditions of elevated global excitation, a discrete buccal oscillator is no longer functionally identifiable.

## Discussion

### Summary of findings

In this study, we present three main findings. We demonstrate that, like the PreBötC in mammals, the Buccal area can endow isolated caudal brainstem segments with endogenous bursting ability when isolated and excited. In the more intact network, we illustrate that a slight increase in network excitation having minor effects on overt motor patterns recruits rhythmically active buccal premotor units beyond the buccal area. This recruitment includes transformation of premotor lung units into buccal units. Finally, we show that under these conditions, the necessity (i.e., essential nature) and sufficiency (i.e., importance) of the buccal area are lost, with the buccal oscillator seemingly replaced by a larger network in which rhythm generation is broadly distributed. Collectively, these results suggest that even when the overt ventilatory motor pattern is invariant, the functional characteristics of premotor units within the frog respiratory network are state dependent. At the network level this state dependence can lead to buccal oscillator expansion causing it to lose its discrete anatomical importance such that rhythm generation becomes distributed across a large population of neurons. To our knowledge, this provides the first evidence that ventilatory rhythm generation in non-mammalian tetrapods can become distributed at the network level beyond a fixed multi-oscillator architecture, raising the possibility that distributed, state-dependent rhythmogenesis represents a conserved principle across vertebrates, including mammals.

### Functional comparison between buccal and PreBötC oscillators

In studying ventilatory rhythm generation in the frog and comparing it with its mammalian counterparts, we have found it important to seek criteria to define both rhythm generating circuit elements in general and oscillators in particular. Accordingly, and based on formative work on invertebrates Central Pattern Generator’s (Marder and Bucher, 2001), we define rhythm generating network elements as elements that pass three criteria: (i) able to increase frequency when stimulated (this has been described as the “sufficiency test”) (Kupfermann and Weiss, 1978); (ii) able to slow or stop a rhythm when the element is functionally ablated (the “necessity test”) (Marder and Calabrese, 1996); and (iii) have activity that is correlated with the rhythm (the “correlation test”) (Marder and Calabrese, 1996). However, from a practical perspective, necessity and sufficiency may not work in a large degenerate network where rhythm generation is distributed across many elements.

In the frog isolated brainstem preparation under normal conditions, the Buccal area passes sufficiency, necessity and correlation tests for being a part of the buccal rhythm generating circuit. Similarly, the region of the ventral lateral medulla in mammals containing the PreBötC passes these tests for inspiration (Alsahafi et al., 2015; Gray et al., 2001; Wang et al., 2014). These tests identify important sites in the rhythm generating circuit, but they do not fully explain the role of these sites in the context of the intact network.

The PreBötC was initially identified using a different test to those above, that of the ability to produce bursts in isolation (Smith et al., 1991). Using isolation as a test for a rhythm generator is not without controversy, largely because motor patterns produced by isolated sections of the nervous system rarely resemble the motor pattern of more intact preparations with regards to burst shape, phase relationship and frequency (St John, 2009). Moreover, isolation focuses attention on elements that are endogenously rhythmic - which we refer here as neuronal oscillators - but the mechanics of endogenous rhythmicity, especially when the rhythm is transformed by isolation, may not reflect the mechanism of rhythm generation in the intact circuit. Notwithstanding these important caveats, both rostral and caudal chunks (and thin slices) of the mammalian brainstem containing r7/8 are capable of generating rhythmic activity from respiratory nerves (Ballanyi and Ruangkittisakul, 2009; Ruangkittisakul et al., 2014). Similarly, we previously reported that transecting the brainstem at the level of CN9 produces a caudal section capable of generating a rhythm resembling that of buccal ventilation (Gdovin et al., 1999; Wilson et al., 2002). In this study, we attempted to systematically isolate the buccal oscillator in a slice. Our results parallel those reported in mammals in that we were able to identify a sub-section of brainstem containing r7/8 capable of endogenous rhythmicity. As has been reported in mammals, as the section of brainstem was reduced along the rostral-caudal axis so too was spontaneous bursting such that tonic exogenous excitation was required to sustain rhythmicity (Del Negro et al., 2002b; Kam et al., 2013). In the frog, when the slice was ∼1.8 mm thick, it had the ability to produce a rhythmic output and bursting resembling the normal buccal rhythm when exposed to 125 nM AMPA. In fact, the thinnest rostral-caudal slice capable of producing rhythmic activity was ∼0.8 mm thick; however reliable bursting in this slice required 125 nM AMPA. Whether this bursting reflects buccal activity was indeterminate because, like rodent preparations, the frequency and burst durations differed considerably to that of the normal ventilatory motor pattern (St John, 2009). Nonetheless, these data suggest that a thin region of the brainstem centered around r7/8 has neuronal oscillator properties in the frog, similar to those that have been identified in rodents and likely originate in the PreBötC (Koshiya and Smith, 1999). These data further support our hypothesis that the Buccal area and PreBötC share a common evolutionary origin (Baghdadwala et al., 2016, 2015).

### Network excitation causes oscillator expansion

The transection experiments revealed that a small section of brainstem in the region of the buccal area is capable of endogenous rhythmicity when isolated and excited. This raises the question as to how oscillators respond to excitation. One possibility is that rhythm generation is confined to a discrete anatomical location, and activity within this location increases with excitation. However, our data supports another possibility, that excitation causes an anatomical expansion of the oscillator implying that oscillators are dynamic in structure. We perform two sets of experiments to determine if the buccal oscillator undergoes anatomical expansion with increased network excitation. First, we conducted the extracellular survey in both the whole brainstem and the buccal segment. Our results demonstrate that, upon increased network excitation, there was a significant increase in the proportion of rhythmically active premotor buccal units, and they were found in an expanded region of brainstem. Additionally, we showed that this increase in the number and proportion of premotor buccal units is due in part to recruitment of sub-threshold buccal neurons that, under low excitability, only fire during lung bursts (i.e., operationally defined as lung units). This observation is congruent with intracellular descriptions of lung neurons in this region (Kogo and Remmers, 1994). Second, we tested the necessity and sufficiency of the Buccal area before and after the brainstem was mildly excited. We found that the Buccal area, defined by necessity and sufficiency before excitation, disappeared with excitation, i.e., after increasing network excitation, microinjection of AMPA and GABA into the same area failed to increase and decrease the frequency of buccal bursts, respectively. In these experiments, the efficacy of GABA was determined by its ability to reverse the effects of local AMPA injection. Care was also taken to ensure that the location and volume of injections were the same. Moreover, the wash in and wash out dynamics of the microinjections were very rapid, indicating that only a small region of the brainstem was being affected. Under these circumstances, bath application of 60 nM AMPA which has a minor effect on overt motor patterns was sufficient to undo the local effects of the 5 µM AMPA and 5 mM GABA. Thus, bath application of 60 nM AMPA likely expands the buccal oscillator to occupy a larger region of the caudal brainstem creating a distributed architecture that makes local perturbations to the original Buccal area inconsequential.

### Distributed Rhythm Generation with Regional Specialization

The idea that respiratory rhythm generation is somewhat distributed is not new. Early studies in elasmobranchs and teleosts revealed that gill ventilation arises from a longitudinal column of neurons extending across the rostral-caudal neural axis (Hyde, 1904; Shelton, 1961; Woldring and Dirken, 1951). Our own work in teleosts further supports this view, showing a graded degradation of rhythm generation with progressive rostral-caudal network inhibition—evidence of a distributed architecture rather than discrete oscillators (Duchcherer et al., 2010). Similarly, early mammalian studies by Lumsden and more recent investigations confirm that both the pons and medulla are necessary for eupneic breathing, and that incremental disruption of the pontomedullary circuit in situ perturbs respiratory rhythm (Lumsden, 1923; Smith et al., 2007). Nonetheless, work in mammals and frogs suggest functionally specialized oscillators within the greater brainstem circuit, such as the buccal and lung area in frogs, and the PreBötC, PiCO and pFRG in mammals (Anderson et al., 2016; Baghdadwala et al., 2015; Smith et al., 2013). Our data in frogs complement and extend the rodent studies of Baertsch et al. (2019) in showing that the functional contribution of classically defined rhythmogenic regions is highly state dependent and emerges from interactions within a broader, distributed network, rather than being rigidly confined to anatomically fixed oscillators. While Baertsch et al. demonstrated that reduced inhibition expands the inspiratory rhythmogenic region in rodents, we show that increased excitation produces an analogous functional expansion in frogs, suggesting that the ability of rhythmogenic networks to reconfigure is robust, regardless of the direction of perturbation. Baertsch et al. showed that tonic or silent neurons could be recruited to become phasically active during inspiration; here, we extend this finding by demonstrating that neurons already assigned to one part of the ventilatory pattern (lung) can be dynamically reassigned to another (buccal). Furthermore, by ablating motor neuron pools, we directly demonstrate that this network reconfiguration, including both recruitment and reassignment, occurs at the level of premotor neurons. Together with the evolutionary distance between amphibians and mammals, these findings suggest that state-dependent redistribution of rhythmogenic capacity is likely a conserved vertebrate principle rather than a rodent specialization.

Strikingly, this architectural flux can occur with only minor changes in overt motor output. Functional reorganization can be subtle yet profound, abolishing the essential nature of previously defined hotspots without altering behavior. These findings challenge the assumption that rhythm-generating regions are fixed, highlighting instead the conditional and context-sensitive contributions of different parts of the network.

This dynamic flexibility is not without precedent. Invertebrate systems have long demonstrated that multiple network configurations can generate the same motor output (Bal et al., 1988; Kristan et al., 2005). Moreover, individual rhythmogenic neurons in these systems can transition between firing states or assume different roles depending on network demands. Our observation of lung-to-buccal neuron transitions in frogs echoes this principle, reinforcing the idea that rhythm-generating networks are not only distributed but composed of flexible, interchangeable cellular components, placing amphibian ventilatory networks among the clearest vertebrate examples of this phenomenon.

While discrete oscillators have been identified in reduced, anesthetized, or hypoxic preparations (Anderson et al., 2016; Del Negro et al., 2009; Hayes et al., 2012; Onimaru et al., 2015, 1989; Onimaru and Homma, 1987; Wang et al., 2014), which are conditions likely to suppress broader network activity, their necessity is often harder to demonstrate in vivo. The size of lesions required to disrupt rhythm in intact animals often exceeds the dimensions of functional nuclei identified in reduced preparations (McKay et al., 2005; McKay and Feldman, 2008; Vann et al., 2018; Yackle et al., 2017) and studies in goats show that even large, potentially lethal ablations can be compensated for when initial life support is provided (Forster et al., 2010; Krause et al., 2009). Recent large-scale recordings in awake or in situ preparations further challenge the discrete oscillator model, consistently failing to reveal discrete anatomical boundaries between respiratory neuron clusters (Dhingra et al., 2020, 2019).

A model in which rhythm generation is anatomically and functionally dynamic, involving a broadly distributed network, need not exclude specialized regions. Distinct areas of the brainstem may contribute preferentially under certain states to different phases or conditions of breathing, explaining the appearance of multiple oscillators in isolated preparations without requiring them to act independently in the intact system. Such architecture confers robustness, adaptability, and resilience, qualities that are essential for maintaining vital rhythmic function in the face of injury, metabolic stress, or developmental abnormalities (Mellen, 2010). For example, in critical conditions, the network’s ability to collapse into minimal core oscillators may serve as a final fail-safe mechanism to sustain life (St John, 2009).

### Technical considerations of method used to functionally ablate motor neurons

To ensure that only premotor units were analyzed, we used a novel approach to functionally ablate specific motor neuron populations. This approach utilized prolonged high-frequency stimulation of cranial nerves applied through extracellular suction electrodes to induce cell death. To limit excitotoxicity to motor neurons, nerve stimulation was performed in high Mg^2+^ to block chemical synapses. On washing out the high Mg^2+^, rhythmic activity returned to all nerves except those that had been stimulated; if only one nerve was stimulated, the contralateral nerve was unaffected and capable of expressing normal ventilatory motor patterns. To the best of our knowledge, this is the first time that this approach has been used to inactivate motor neurons to investigate rhythm generating circuits. Nevertheless, high frequency stimulation is commonly used to simulate the effect of seizures and epilepsy – both of which result in cell death -- and nerve stimulation has also been demonstrated to cause motor neuron death in chick embryos (Kienzler et al., 2006; Olney et al., 1983; Oppenheim and Núñez, 1982).

We carried out a series of experiments to validate the prolonged high-frequency stimulation approach in the frog. To confirm efficacy of our methods, we conducted time control experiments to ensure that motor neurons were functionally ablated for the entire duration of our protocol (∼4 h) and not just temporarily silenced. To further confirm that the motor neurons were functionally ablated, we performed pilot experiments involving brief stimulation of one nerve to stimulate reflex responses in others. Brief stimulation of nerves subjected to prolonged high-frequency stimulation, failed to induce activity in healthy nerves; and brief stimulation of healthy nerves, while sufficient to induce activity in other healthy nerves, failed to induce responses in nerves subjected to prolonged high-frequency stimulation (data not shown). Of note, when contralateral nerves spanning the buccal area (i.e., left and right CN10 and SN1; CN9 is not respiratory in the frog) were subjected to prolonged high-frequency stimulation, CN5 continued to express both buccal and lung bursts when high Mg^2+^ was washed out, indicating that caudal brainstem motor neurons are not a necessary part of the buccal oscillator.

### Limitations and future directions

Our findings are entirely focused on the Buccal area. Further studies are needed to determine if the lung and priming areas are similarly state dependent. In addition, while the isolated frog brainstem produces rhythms closely resembling those in vivo, the network architecture and its ability to reconfigure might be quite different in the presence of descending and sensory input. Another limitation is that our work is solely on the frog; we often argue that the fundamental architecture of the rhythm generating circuit is conserved across vertebrates, however, this remains conjecture until more grounded data demonstrating evolutionary lineage of respiratory oscillators is available.

## Abbreviations

aCSF: artificial Cerebral Spinal Fluid
CN: cranial nerve
IBI: inter buccal interval
IF: instantaneous frequency
preBötC: preBötzinger Complex
SD: standard deviation
SN: spinal nerve

## Acknowledgments

This work was funded by a Discovery Grant from National Science and Engineering Council (NSERC/ RGPIN-2020-05312) of Canada. Salary support for R.J.A.W was provided by Alberta Innovates Health Solutions (AIHS). M.I.B was supported by a Summer Studentship from the O’Brien Centre, a Gerald L. Weber Scholarship and an NSERC Studentship. M.R.S. was supported by a CIHR Postdoctoral Fellowship (REH 187964), SCNIP and Brain CREATE awards.

## Author contributions

Conceptualization of study and methodology devised by M.I.B and R.J.A.W. Experiments conducted, data curated, analyzed and visualized, and manuscript written by M.I.B., M.R.S and R.J.A.W. Operating funding obtained by R.J.A.W. All the authors approved the final version of the manuscript submitted for publication. All authors agree to be accountable for all aspects of the work. All people designated as authors qualify for authorship, and all those who qualify for authorship are listed.

